# Life Strategies in Placozoa

**DOI:** 10.1101/2021.11.26.470175

**Authors:** Daria Y. Romanova, Mikhail A. Nikitin, Sergey V. Shchenkov, Leonid L. Moroz

## Abstract

Placozoans are essential reference species for understanding the origins and evolution of the animal organization. However, little is known about their life strategies in natural habitats. Here, by establishing long-term culturing for four species of *Trichoplax* and *Hoilungia*, we extend our knowledge about feeding and reproductive adaptations relevant to their ecology and immune mechanisms. Three modes of population growth depended upon feeding sources, including induction of social behaviors and different reproductive strategies. In addition to fission, representatives of all haplotypes produced ‘swarmers,’ which could be formed from the lower epithelium (with greater cell-type diversity) as a separate asexual reproduction stage. In aging culture, we reported the formation of specialized structures (‘spheres’) from the upper cell layer as a part of the innate immune defense response with the involvement of fiber cells. Finally, we showed that regeneration could be a part of the adaptive reproductive strategies in placozoans and a unique model for regenerative biology in general.

## INTRODUCTION

Placozoans are essential reference species to understand the origins and evolution of the animal organization. Despite the long history of investigations, Placozoa is still one of the most enigmatic animal phyla. Placozoans have the simplest known animal body plan – three cell layer’s organization (Schultze, 1883; Metschnikoff, 1886, 1892; Noll, 1890; Graff, 1891; Stiasny, 1903; Ivanov, 1973; Rassat and Ruthmann, 1979; Dogel, 1981; Malakhov and Nezlin, 1983; Okstein, 1987; Malakhov, 1990; Smith et al., 2014; Romanova, 2019; Mayorova et al., 2019; Smith et al., 2019; Romanova et al., 2021), but surprisingly complex behaviors (Kuhl and Kuhl, 1963, 1966; Seravin and Karpenko, 1987; Seravin and Gudkov, 2005; Eitel and Schierwater, 2010; Eitel et al., 2013; Smith et al., 2015; Senatore et al., 2017; Armon et al., 2018; Cuervo-González et al., 2020) with social feeding patterns (Okstein, 1987; Fortunato and Aktipis, 2019).

The phylum Placozoa contains many cryptic species. Differences in phenotypes are minor and sampling across the globe revealed about 30 haplotypes (Miyazawa et al., 2012; Eitel et al., 2010, 2013; Aleshin et al., 2004; Miyazawa et al., 2021; Schierwater and DeSalle, 2018), but there are only three formally described species: *Trichoplax adhaerens* (Schultze, 1883), *Hoilungia hongkongensis* (Eitel et al., 2018) and *Polyplacotoma mediterranea* (Osigus et al., 2019).

Placozoans were collected predominantly from tropical and subtropical regions (Ueda et al., 1999; Signorovitch et al., 2006; Pearse, 2007; Eitel and Schierwater, 2010; Nakano, 2014); they could live in a wide range of salinity (20-55 ppm), temperature (11-27°C), depth (0-20 m), and pH (Schierwater, 2005; Schierwater et al., 2010; Eitel et al., 2013).

However, their lifestyle is essentially unknown in nature. Pearse (2007) had suggested that placozoans may be opportunistic grazers, scavenging on organic detritus, algae, and bacteria biofilms. Long-term culturing helps to explore the life history of placozoans further. Most of the knowledge about placozoans had been obtained from culturing of just one species, *Trichoplax adhaerens* (Grell strain, H1 haplotype, see details in (Signorovitch et al., 2006; Eitel and Schierwater, 2010; Eitel et al., 2013; Heyland et al., 2014; Pearse, 1989)). Both rice and algae had been used as alternative food sources (e.g., *Cryptomonas* (Grell, 1972, Ruthman, 1977), red algae *Pyrenomonas helgolandii* (Signorovitch et al., 2006), green algae (*Ulva sp*; Seravin, Gerasimova, 1998), or a mix of green, *Nannochloropsis salina*, and red algae, *Rhodamonas salina, Pyrenomonas helgolandii*, (Jackson and Buss, 2009; Smith et al., 2014)) as well as yeast extracts (Ueda et al., 1999).

Here, by establishing long-term culturing for four species of *Trichoplax* and *Hoilungia*, we provided additional details about feeding and reproductive adaptations relevant to placozoan ecology and immune mechanisms.

## MATERIAL AND METHODS

### Culturing of placozoan haplotypes

We used clonal cultures of four species of Placozoa: *Trichoplax adhaerens* (Grell’s strain H1, from the Red Sea), *Trichoplax* sp. (H2 haplotype, collected in the vicinity of Bali island), *Hoilungia* sp. (H4 haplotype, collected in coastal waters of Indonesia), and *Hoilungia hongkongensis* (H13 haplotype, found in coastal waters of Hong Kong). We maintained all species in culture for 2-3 years that allowed long-term observations and adjustments of culture conditions for each haplotype.

We maintained three haplotypes (H1, H2, and H13) in closed Petri dishes with artificial seawater (ASW, 35 ppm, pH 7.6-8.2), which was changed (70% of the total volume) every 7-10 days. At the same time, a suspension of the green alga *Tetraselmis marina* was added into the culture dishes. Alternatively, we use 5-7 rice grains for 1 Petri dish. When the biofilm of microalgae became thinner or depleted, freshly prepared, 1-2 mL suspension of *T. marina* could be added to the culture dishes. Mixtures of other algal clonal strains were occasionally used as food sources (for example, the cyanobacteria *Leptolyngbya ectocarpi* and *Spirulina versicolor*, the red algae such *Nannochloropsis salina* or *Rhodomonas salina*). We maintain three placozoan haplotypes at the constant temperature of 24 °C and natural light in environmental chambers (**Modes 1-3**). In parallel, H1, H2, and H13 haplotypes were also successfully cultured using rice grains as nutrients (**Fig. 1**).

**Figure 1.**
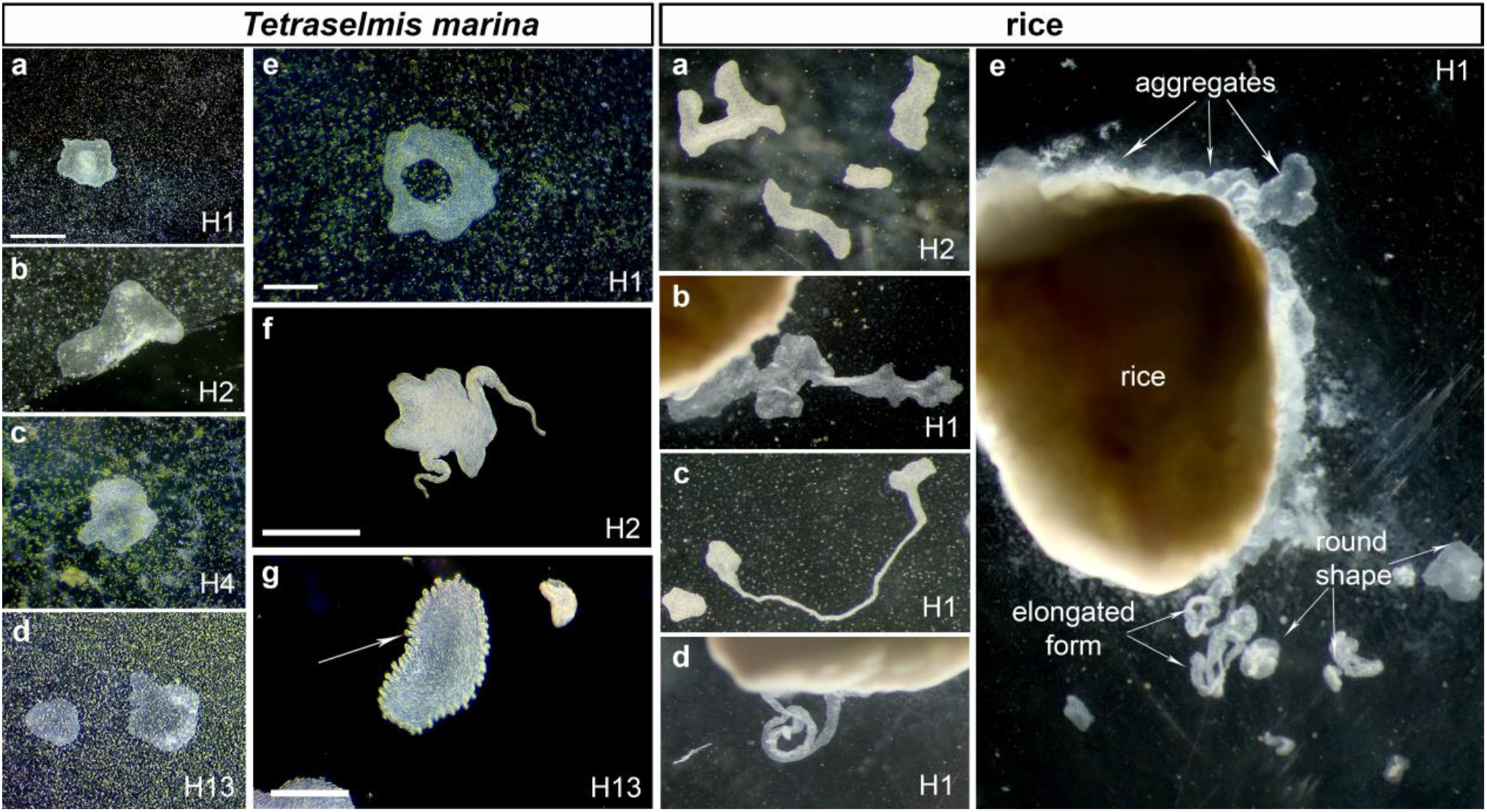
The diversity of placozoan body forms. Illustrated examples from long-term culturing of different haplotypes (indicated in each photo) on two feeding substrates (*Tetraselmis marina* and Rice boxes). *Tetraselmis marina* box shows both canonical placozoans body shapes (a-d) and unusual morphologies (animals with a ‘hole’ in the middle of the body (**e** and supplementary video 1), elongated ‘pseudopodia’-like structures (**Figure S3**, Supplement), and animals with numerous small ovoid formations in the rim area (**g**). Rice grains support a stable population growth rate but often facilitate the formation of aggregations of animals around grains (Rice box). Scale bar: *Tetraselmis marina* box: 500 µm; Rice box: grain: 1 mm.

Placozoans are transparent, but their color could be changed depending on the algae they are feeding on. For example, light-brownish color occurs with *T. marina* as a food source, medium brownish coloration was observed in animals fed on diatoms (*Entomoneis paludosa, Psammodictyon* sp.), or pinkish colors were seen when animals were fed on cyanobacteria. Pinkish coloration might be due to the accumulation of phycobilins from cyanobacteria, red algae, and cryptophytes.

Under long-term culturing, animals divided every 1-2 days without signs of sexual reproduction (Malakhov, 1990; Zuccolotto-Arellano and Cuervo-González, 2020). We observed exponential growth rates for H1 (Avg = 477), H2 (Avg = 312), and H13 (Avg = 232) haplotypes in triplicate experimental groups.

In contrast to other placozoans, the H4 haplotype could be successfully cultured at 28 °C using a mixture of the green algae *T. marina* and two cyanobacteria *Spirulina versicolor* and *Leptolyngbya ectocarpi* (see also Okstein’s, 1987). However, if the H4 haplotype was maintained on *T. marina*, the population growth was significantly declined.

### Cryo fixation for transmission electron microscopy

Animals were placed in cryo capsules 100 µm deep and 6 mm in diameter in ASW. After specimens were adhered to the bottom of capsules, the ASW was replaced with 20% bovine serum albumin solution (in ASW). The animals were fixed with a High-Pressure Freezer for Cryofixation (Leica EM HPM 100 for cryofixation of Biological and Industrial Samples). After fixation, animals were embedded in epoxy resin (EMS, Hatfield, UK). Ultrathin (65 nm) serial sections were made using Leica EM UC7 ultramicrotome. Sections were stained in uranyl acetate and lead citrate (Reynolds, 1963) and studied using JEOL JEM-2100 and JEOL JEM-1400 (JEOL Ltd., Tokyo, Japan) transmission electron microscopes with Gatan Ultrascan 4000 (Gatan Inc., Pleasan-ton, CA) and Olympus-SIS Veleta (Olympus Soft Imaging Solutions, Hamburg, Germany) TEM cameras. TEM studies were done at the Research center “Molecular and Cell Technologies” (Saint Petersburg State University).

### Laser scanning microscopy

Swarmers and ‘spheres’ were transferred from cultivation Petri dishes to sterile Petri dishes using a glass Pasteur pipette. Individual animals were allowed to settle on the bottom of the container overnight. Fixation was achieved by gently adding 4% paraformaldehyde in 3.5% Red Sea salt (at room-temperature) and stored at 4 °C for 1 hour. We washed animals in 1% of bovine serum albumin, then incubated them for 15 min in Mito-tracker Green. Next, preparations were washed in phosphate buffer solution (0.1 M, pH=7.4, 1% Tween-20) three times (20 min) and mounted on a slide using Prolong gold antifade reagent with DAPI, and stored in the dark at 4°C.

The samples were examined using Zeiss LSM 710 confocal laser scanning microscope with a Plan-Apochromat 63x/1.40 Oil DIC M27 immersion lens (Zeiss, Germany). The images were obtained using the ZEN software package (black and blue edition) (Zeiss, Germany). Image processing was carried out using ZEN (blue edition), Imaris, ImageJ software.

### Statistical analysis

Population growth rate, locomotion, regeneration analysis and occurrences of aggregates were calculated using standard statistical (t-test) and heatmap packages in R. We use triplicates for population growth rates; see additional details in the Result section.

### Treatment of *T. adhaerens* with antibiotics

*Trichoplax* might contain potentially symbiotic bacteria in fiber cells (Driscoll et al., 2013; Kamm et al., 2019). To control levels of bacterial endosymbionts, we used treatment with different antibiotics (ampicillin (5 µg/mL), doxycycline (1.25 µg/mL), ciprofloxacin (7 µg/mL), and rifampicin (1.25 µg/mL).

Total DNA from individual animals was extracted using a silica-based DiaTom DNAprep 100 kit (Isogene, Moscow, Russia) according to manufacturer’s protocol. Amplification was performed using EncycloPlus PCR kit (Evrogen, Moscow, Russia) using the following program: 95 °C – 3 min, 35 cycles of PCR (95 °C – 20s, 50 °C – 20s, 72 °C – 1 min), and 72 °C – 5 min. We have used universal forward primer 27F (AGA GTT TGA TCM TGG CTC AG) and specific reverse primer 449R (ACC GTC ATT ATC TTC YCC AC). The reverse primer was designed against 16S RNA of *Rickettsia belli* (NR_074484.2) and sequences from *Trichoplax* DNA found through NCBI Trace Archive Blast using NR_074484.2 as query. After 6 months of ampicillin treatment the *Rickettsia* were not detected. Other antibiotics were less effective (Fig. 5S, see Supplement).

## RESULTS

### 1. Three states of long-term culturing and feeding in Placozoa

Analysis of growth and behavioral patterns during the long-term culturing allowed us to distinguish three distinct conditions, which shared across placozoans.

#### Optimized culture conditions

This first mode describes a population with a stable growth rate on the established algal mat (∼6.5×10^6^ cells of *T. marina* per 1 μL, added once a week) or rice grains (4-5 grains in one Petri dish, diameter 9 cm), and refreshing liquid medium once in 7-10 days. We observed regular fission of placozoans, at average once a day, during a few months (**Fig. 3**, Mode 1). Thus, we transferred an excess of animals to other dishes maintaining about 500 individuals per dish. This culturing allows keeping stable populations of placozoans for a long time (from a few months to 2-5 years).

**Figs. 1-2** show a diversity of canonical placozoans body shapes (**Fig. 1a-d**). However, unusual morphologies were also observed in all haplotypes. For example, we noted animals with a ‘hole’ in the middle of the body (**Fig. 1e**,), elongated ‘pseudopodia’-like structures (**Fig. 3S, Supplement**), or numerous small ovoid formations in the rim area (**Fig. 1g**).

**Figure 2.**
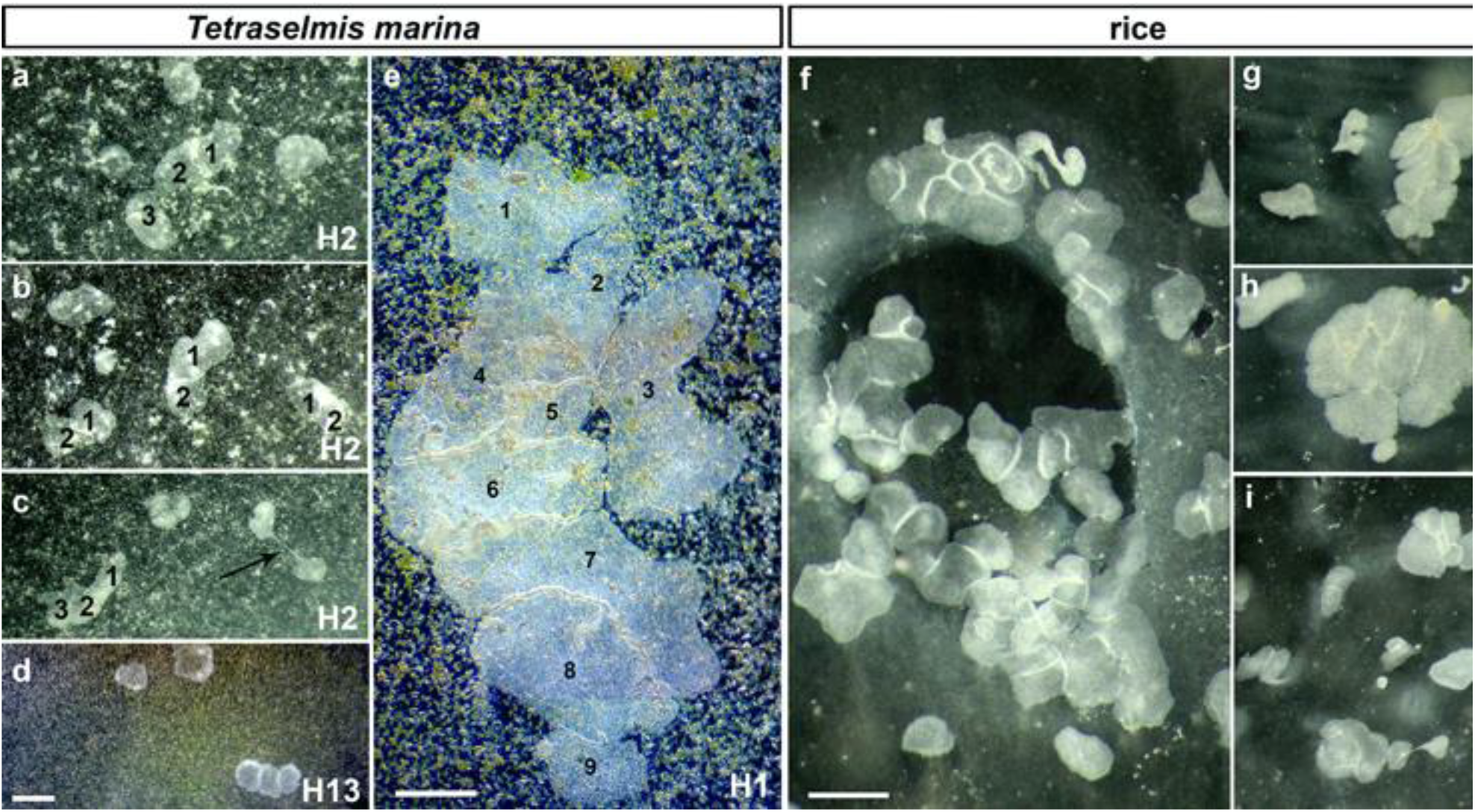
Social feeding behavior during long-term culturing on the dense algal. (*Tetraselmis marina* box) **and rice substrates.** The aggregation of animals depended upon the density of the substrates (see mode #3 in the text and Fig. 3). This behavioral pattern was observed for all haplotypes (H1, H2, H13 – in Petri dishes, H4 – in 20L aquaria). Aggregates often included 2-15 individuals (e.g., 9 H1 individuals in e). pH=8.2. Scale bar: a-d – 100 µm, e – 1 mm, f-i – 500 µm.

**Figure 3.**
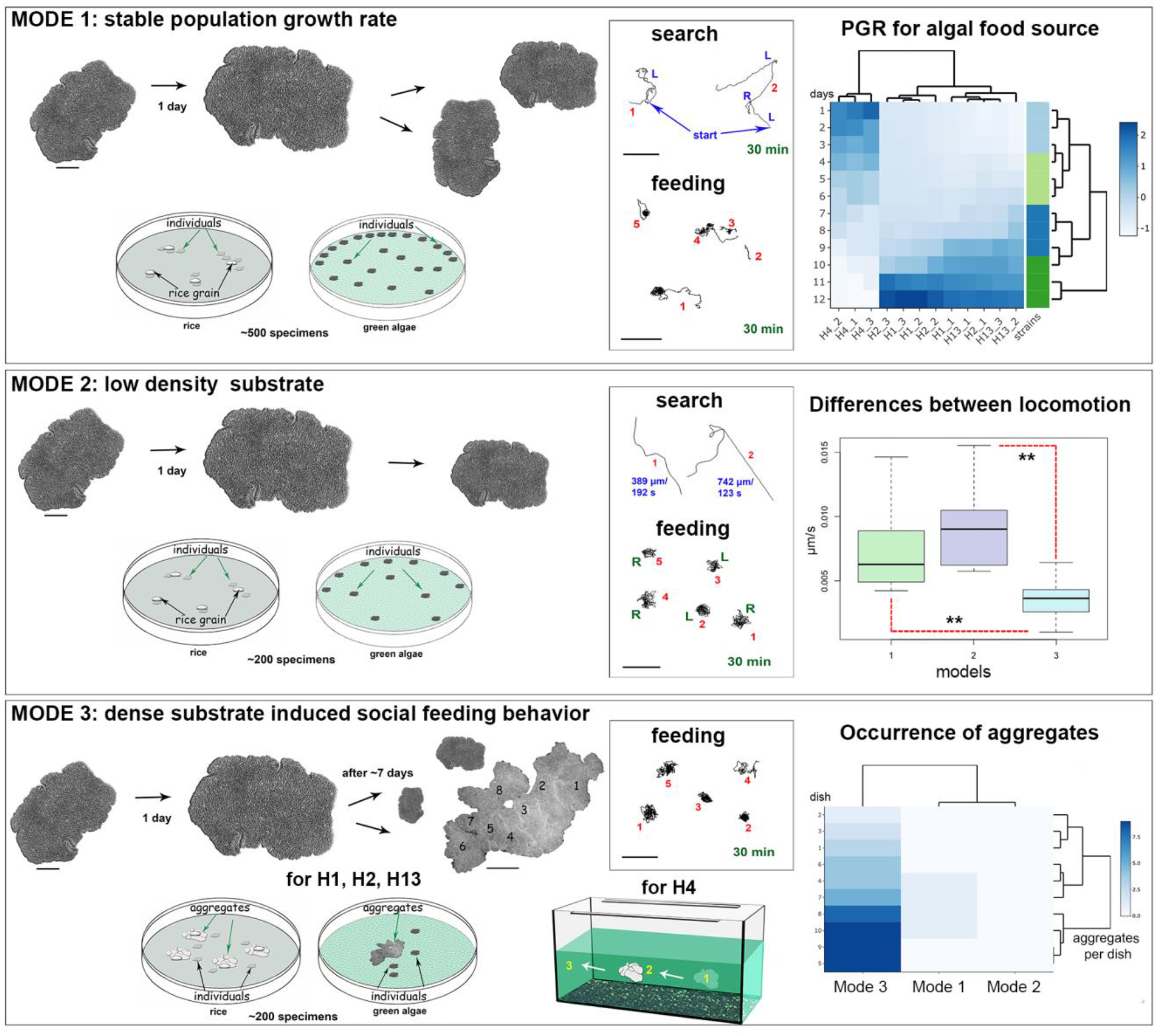
Three separate model schemes of long-term culturing/aging of Placozoa. Mode 1 – optimized conditions; Mode 2 - low and Mode 3 - high densities of the green alga mat (Tetraselmis marina) or rice grains; see text for details. Mode 1: Optimized culture conditions with dispersal animals and their moderate concentration on feeding substrates. The middle diagram shows examples of exploratory and feeding locomotion for five animals (numbers) within 30 min. The right diagram shows the representative dynamics of the population growth rate (PGR) for all haplotypes (H1, H2, H4, and H13). 3 separate dishes were used for each haplotype (e.g., H4_1, H4_2, H4_3, etc.), starting with 10 animals per dish. All datasets were normalized to absorb the variation between columns for all four haplotypes of Placozoa. Under these conditions, animals steadily increase their body surface area and have asexual reproduction by fission. Mode 2: Low-density substrate. Limit of food source led to decreasing of animal sizes and numbers of animals in culture dishes. Mode 3: High-density substrate. There is both increasing in animal sizes and the aggregation of 2-15 individuals around rice grains or on the dense algal mat. The heat diagram on the right shows the predominant occurrence of aggregates compared to Mode 1 (no aggregates were observed on low-density substrates in Mode 2). H4 expressed the same behavior patterns on the walls of 20L aquarium, where individuals within the aggregate could move together (1-2-3, arrows). However, most animals stay at the substrate (central diagram) with significantly reduced overall locomotion during the feeding, as indicated in the right middle diagram. Difference between locomotion in Modes 1 and 3: p-value is 0.003002; between Modes 2 and 3: p- value is 0.000072 (unpaired Student’s test). Scale bars: for individual animals - 200 µm; for the aggregate in Mode 3 – 1 mm; for all locomotory tracks - 200 µm.

#### Depleted food substrate

If no extra food source was added within 2-3 weeks, the biofilm of microalgae became thinner or depleted. When the layer of microalgae became less than 4.2×10^5^ cells/μL, we observed a 1.5-2 fold reduction in animals’ surface areas in all tested haplotypes (H1, H2, and H13), and the population size decreased from ∼500 to ∼200 animals per one cultivation dish (**Fig 3, Mode 2**). Under these conditions, the animals were concentrated in the densest areas of the algal substrate. In 4-5 weeks, several percent of placozoans formed unusual spherical structures described in section 3.

#### High density of food substrate and ‘social’ behavior

The third mode of culturing was observed on dense substrates such as 3–4-layer algal biofilm with 8×10^8^ cell/µL of suspension or 7-8 rice grains (per Petri dish, diameter 9 cm for H1, H2, and H13). Within a few days on abundant food sources, placozoans often formed clusters consisting of multiple animals (**Fig. 2**). These aggregates of 2-15 animals have been described as a type of ‘social’ behavior (Okstein, 1987, Fortunato and Aktipis, 2019). These conditions also induced social feeding patterns in the 20 L aquaria system for the H4 haplotype (**Fig. 3**, Mode 3).

This type of collective behavior differed from typical alterations of search/exploratory and feeding cycles observed in sparked individuals under conditions with limited food sources (Modes 1 and 2, **Fig. 3**). When animals were feeding, they usually stayed on the food substrate or rotated for ∼15-30 mins within small region, comparable to their body length (**Fig. 3**).

### 2. Life strategies

#### 2.1 Reproduction: forming ‘swarmers.’

Long-term culturing provided additional insights into the life-history strategies of Placozoa. In addition to the fission, the formation of smaller daughter animals or ‘swarmers’ had been described in *Trichoplax adhaerens*, and swarmers were reportedly derived from the upper epithelium (Thiemann and Ruthmann, 1990, 1991). Here, we observed the development of swarmers in all haplotypes studied (H1, H2, H4, and H13; **Figs. 4-5**), suggesting that it is an essential part of adaptive strategies for Placozoa (see supplementary **Video 4** for *Trichoplax* sp. (H2), and **Videos 5-6** for *Hoilungia* (H4 haplotypes)).

**Figure 4.**
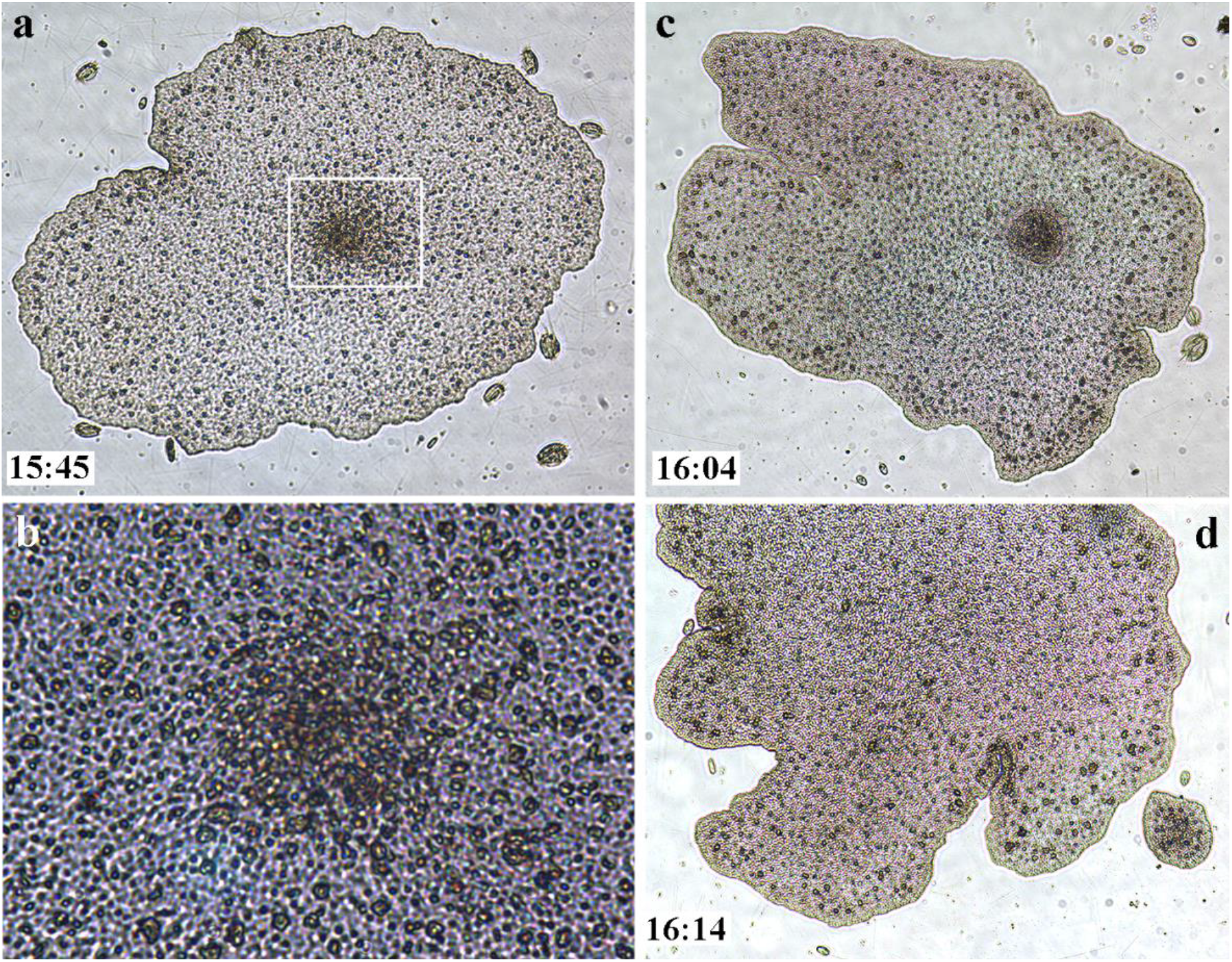
Swarmers were formed at the lower (substrate-facing) side of the ’mother’-animal (H1 haplotype, *T. adhaerens*). a, c, d – A unique illustrated example of the time course of the swarmer formation (bottom left corners indicate time intervals). **a**. The formation of a higher density cell region in the middle part of the mother animal (white square outline, and the higher magnification of the same region in **b**). **c**. The formation of the swarmer and its separation from the mother animal (d).

**Figure 5.**
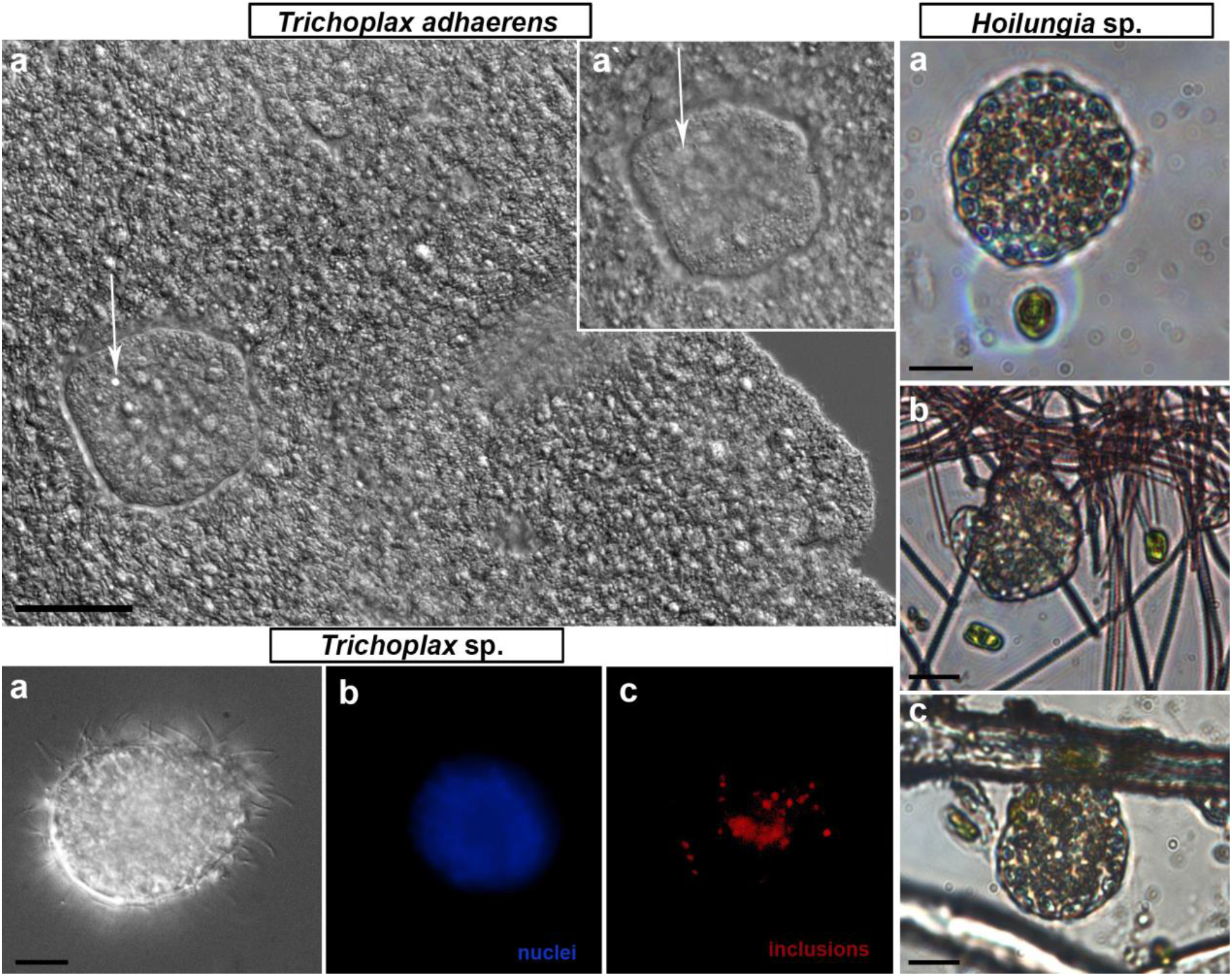
Swarmers were formed in the long-term culture in every studied species of Placozoa. Swarmers are small (15-30 µm diameter) juvenile animals with presumably six cell types: upper and lower epithelial, gland and lipophil cells, fiber, and crystal cells (white arrows in *T. adhaerens* box, a- a’ - different Z-layer). Swarmers possessed the canonical placozoan bodyplan (*Trichoplax* sp. box). Despite their small size, swarmers expressed coordinated exploratory and feeding behaviors (*Hoilungia* sp. box). Scale bar: 10 µm.

The formation of ‘swarmers’ occurred spontaneously (in about 2 or 5 weeks from a cultivation start) both on algal biofilms and rice. But we noted that swarmers could be formed at the lower, substrate-facing side (**Fig. 4**, Supplementary **Video 3**), with a significantly greater cell-type diversity than in the upper layer (Smith et al., 2014, Mayorova et al, 2019, Romanova et al, 2021). Therefore, the formation of swarmers from the lower layer might be facilitated by the preexisting heterogeneity of cell types in this region. After physical separation of ‘swarmers,’ these young animals could be temporally located under the ‘mother’ (supplementary **Video 3**), moving together on substrates or biofilms.

#### 2.2 Regeneration as a part of adaptive life strategies in placozoans

An overpopulated/fast-growing culture with a high density of placozoans (over 500-700 animals per 1 Petri dish 9 cm in diameter) often contains many floating individuals or individuals on walls, which are frequently aggregated under the surface film (**Fig. 2S, supplement**). As a result of contact with air and/or other types of mechanical damage, the animals could be raptured. Still, such fragmentation often led to regeneration, which we consider an essential part of life strategy in placozoans.

We investigated the regeneration in model experiments (using H1 and H2 haplotypes,1-2 mm in size). Individuals were damaged in two ways: mechanical injury by pipetting and by cutting animals with a scalpel (into two parts). The former protocol allows obtaining small cell aggregates (∼20-30 cells) placed in Petri dishes with biofilms of *Tetraselmis marina*. The regeneration process lasted approximately 7-10 days. The first stage of recovery was an increase in cell numbers within the aggregates, which were immobile (**Fig. 6**). Notably, intact placozoans had a negative phototaxis (animals moved to the darkened areas of experimental Petri dishes, see supplementary **Fig. 1S**). However, at the early stages of regeneration, *Trichoplax* aggregates did not respond to changes in light intensity, remaining motionless. After 4^th^ day, locomotion was restored (**Fig. 6**).

**Figure 6.**
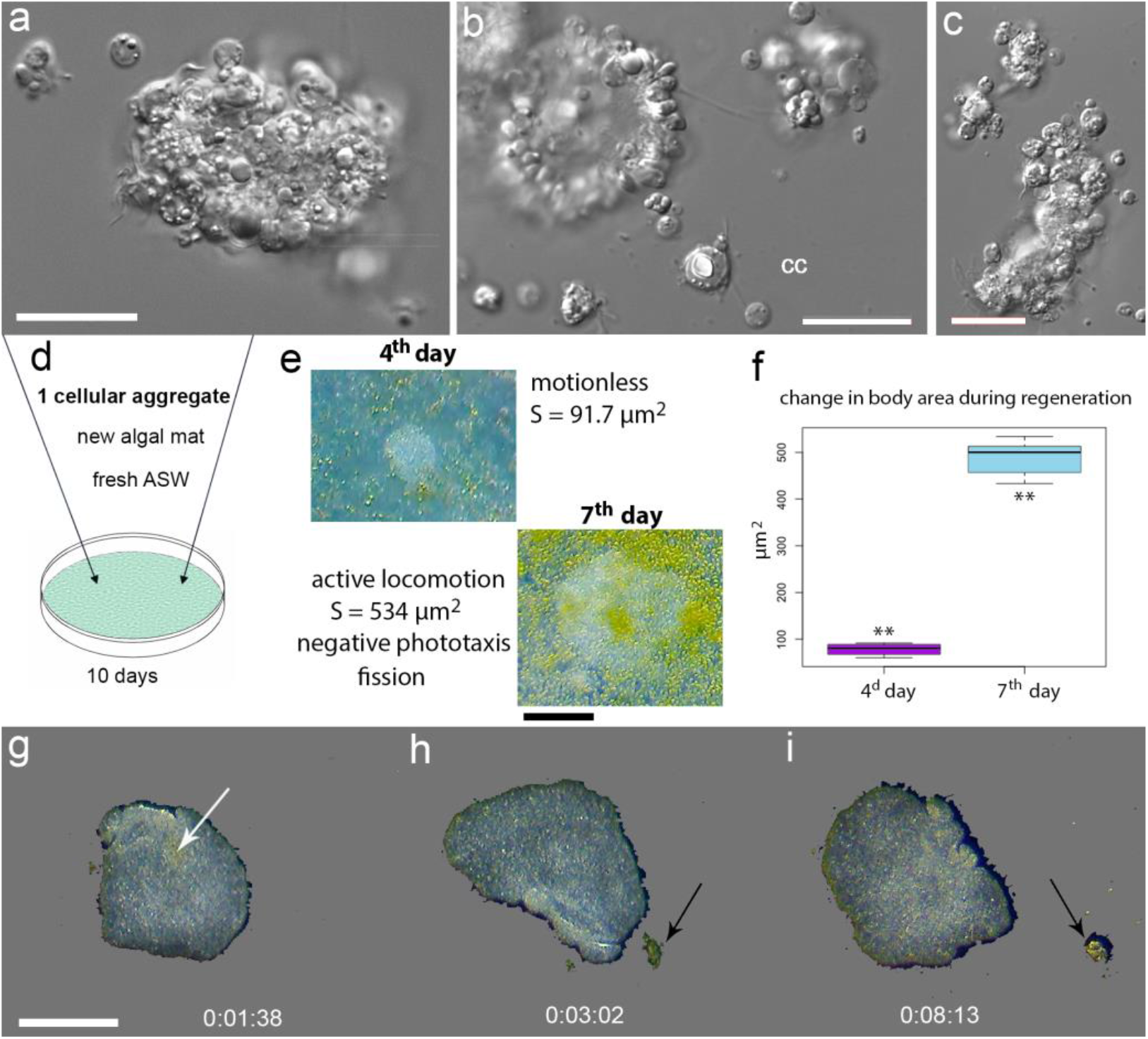
Regeneration in Placozoa. **a, b, c** – Illustrative examples of small cellular aggregates from T. adhaerens. Aggregates consist of ciliated epithelial cells, lipophil, crystal, and fiber cells. cc- isolated crystal cell. **d** - Placement of a single aggregate in a new culture cell with fresh ASW and algal mat. **E** – 4th and 7th day of regeneration with calculated surface area (S) of a newly formed animal. The ciliated locomotion, negative phototaxis and fission started on the 7th day of regeneration. **f** – increasing surface areas occurred from 4th to 7th day. The t-value: -23.48979. The p-value: < 0.00001. **g-i** –After the splitting, an animal into two parts, locomotion and feeding continued (see text and Supplementary video 7). Arrows indicate a cluster of algae. Scale bar: a, b, c - 20 µm, e - 200 µm, g-I – 500 µm.

On 7^th^ day original aggregates became small individual animals with active locomotion and feeding behaviors as well as capable of fission and negative phototaxis. Interestingly, if dissociated cells and aggregates were transferred to Petri dishes without a food source, then after 2-3 days, aggregates were lysed.

After dissection individuals into two parts, we observed a slight contraction of animals. Still, within a few minutes, animals curled up, closing the wound, and continued to move without detectable changes in their locomotion patterns (**Fig. 6**, supplementary **Video** 7).

### 3. Spherical formations & Systemic immune response

In 4-5 weeks (Mode 2 of culturing), with depleting food sources, some animals start to develop specific structures, which we call ‘spheres’ (**Figs. 7-8**). The formation of spheres occurred randomly in 2-5% of individuals, and data reported below are based on observations of about one hundred animals with these structures. We view spheres as a component of adaptive immune (e.g., bacterial infection) responses and separated three stages of this process.

**Figure 7.**
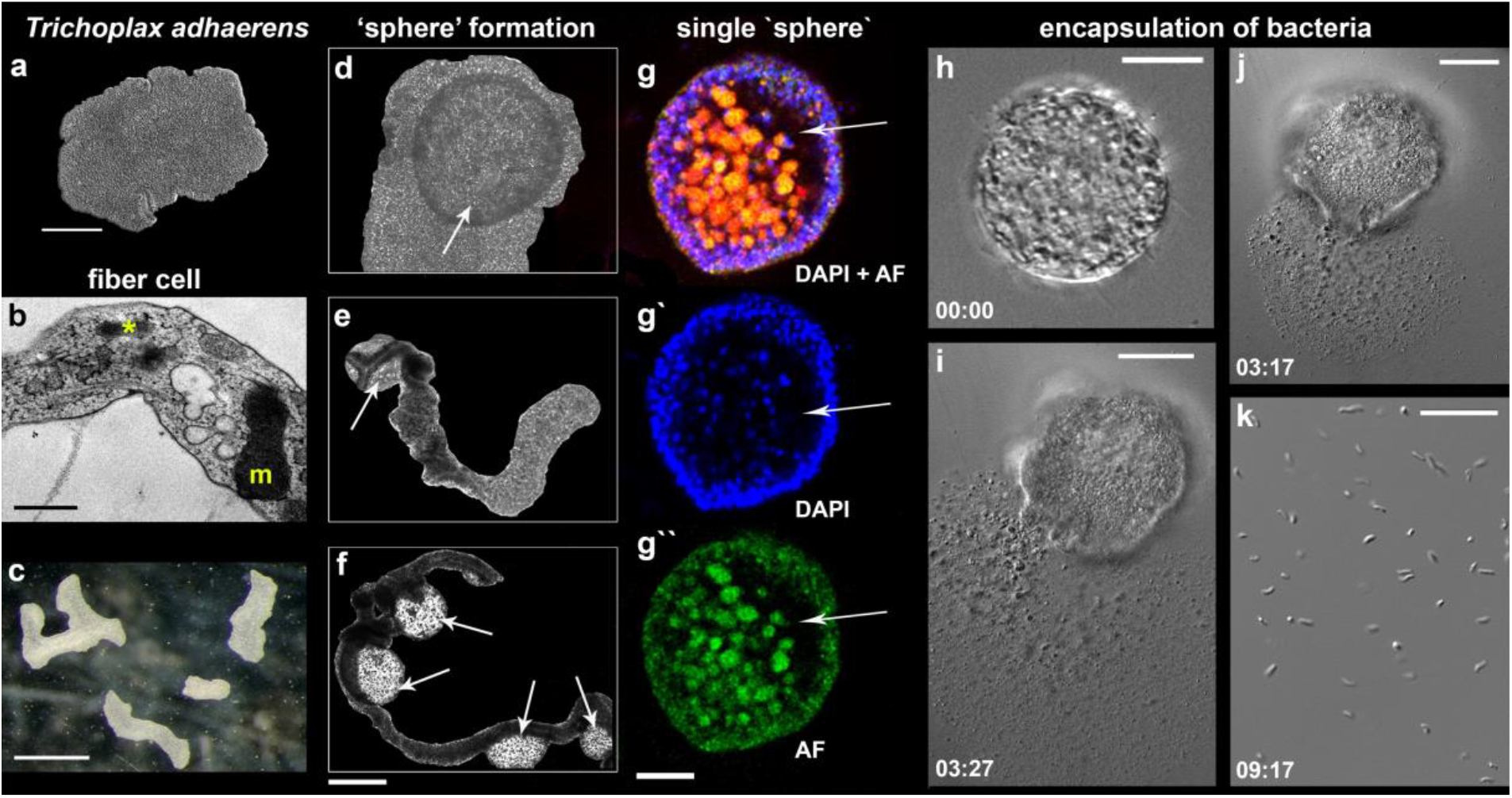
‘Sphere’-type formations in *Trichoplax adhaerens*. **a-c** control animals under optimal culture conditions (mode #1, see text and Fig. 3); **b** – Transmission electron microscopy (TEM) image of the fiber cell with a bacterium (asterisk) and a large mitochondrial complex (m). **d-h**– Formation of spheres (arrows). **g-g’’** – Separated spheres with internal cavities (arrows); Nuclear DAPI staining – blue (excitation 660 nm and emission 682 nm); Autofluorescence (AF) – green (excitation 490 nm and emission 516 nm). **j-k** - Spherical formations encapsulate bacteria inside; the damage of the sphere’s surface by laser released numerous bacteria (see text for details). Time intervals following the laser-induced injury are indicated in the left corners of each image. Scale: a 200 µm, b – 500 nm, c – 1 mm, g-g’’ - 400 µm, g-i - 20 µm, k - 10 µm.

**Figure 8.**
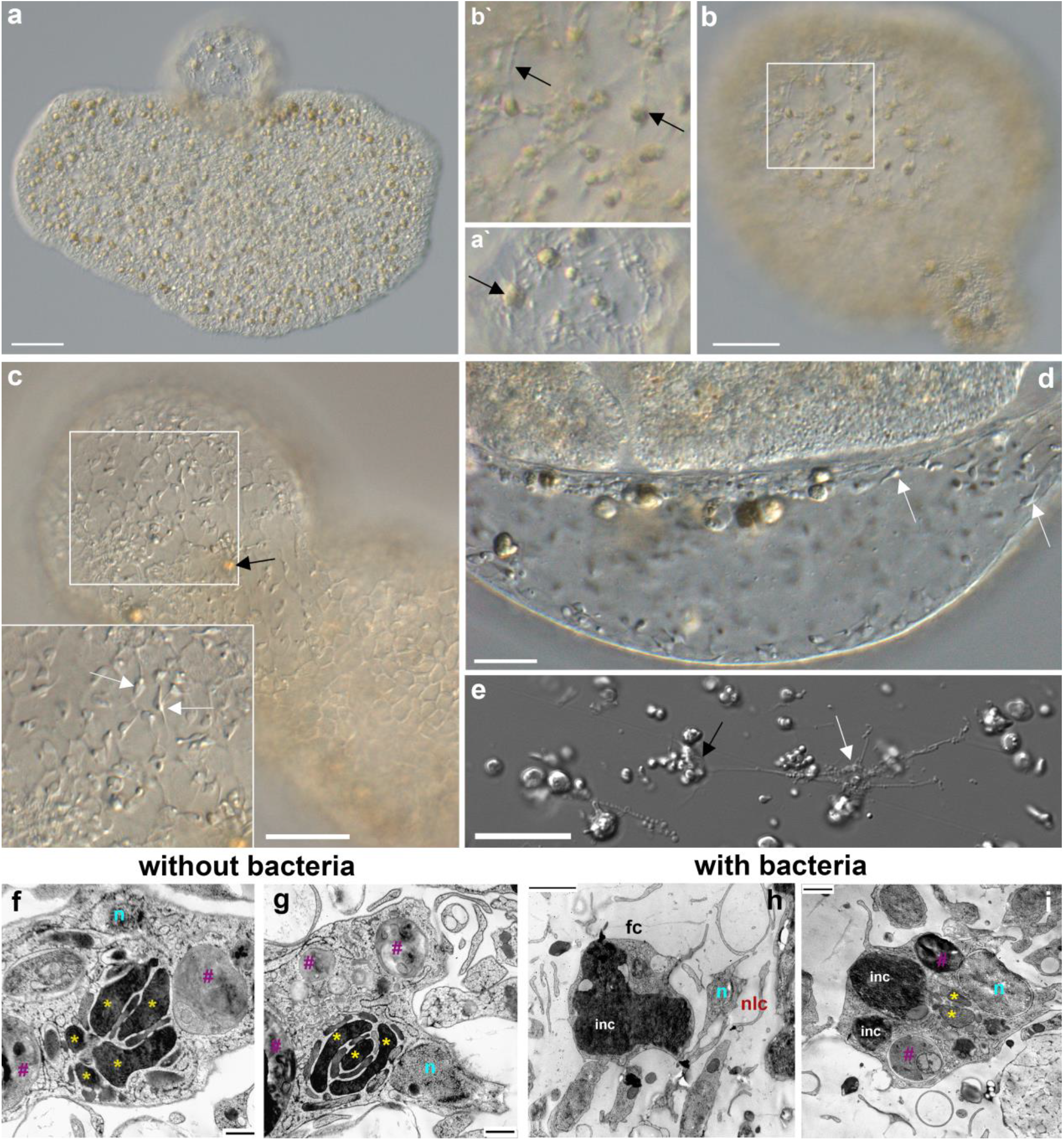
Fiber cells in spheres (a-c) and their ultrastructure (f-i). Both fiber cells (black arrows) and smaller neuroid-like cells (white arrows) are present in ‘spheres’. a - *Trichoplax adhaerens* with a sphere on the upper side. a’– a part of the sphere with two fiber cells (arrow). c, d, e – smaller neuroid- like cells (white arrows) with elongated processes (e) that can form connections among themselves and fiber (black arrow) cells (Romanova et al., 2021). Both cell types are located in the middle layer of placozoans (b/b’). f-i – Transmission electron microscopy (TEM) of the fiber cells in *Rickettsia*-free *Trichoplax* (ampicillin-treated for 12 months) and control animals with endosymbiotic bacteria (see also Fig. 7b). Of note, fiber cells in ampicillin-treated populations of placozoans had more elaborate mitochondrial clusters and clear inclusions. In contrast, in animals with bacteria, fiber cells possessed large dark (by TEM) inclusions (light microscopy also shows brownish inclusions – black arrows in a,b,c). Abbreviations: white arrows – neuroid-like cells, black arrows – fiber cells, fc – fiber cell, nlc – neuroid-like cell, inc – inclusion, yellow stars – mitochondria, n – nucleus, purple pad key – other inclusions. Scale bar: a-c, e – 20 µm; d – 10 µm; f, g – 500 nm, h-i – 1 µm.

The first stage was an apparent lengthening of body shape (**Fig. 7**c) and inability to fission (**Fig. 7**c-f). This phenomenon was observed during aging of populations, which occurred without systematic refreshing of ASW. When ASW was replaced, within 2-3 days, we had restored populations without any noticeable morphological changes compared to control individuals (as in Mode 1).

The second stage: the lower epithelium begins to exfoliate (**Fig. 7**d-f), and spherical formations become visible. We could observe elongated animals with one ‘sphere’ (**Fig. 7**e) and several spheres (**Fig. 7**f), as well as rounded animals with one ‘sphere’ (**Fig. 7**d).

The third stage was a separation of the ‘spheres’ from a main body (**Fig. 7**g-g’’). ‘Spheres’ consisted of lower epithelium, fiber cells, and, probably, ‘shiny spheres’ cells contained large lipophilic inclusions. Ultrastructural analysis showed cavities inside ‘spheres’ (**Fig. 7**g-g’’) and large autofluorescent granules of unknown etiology. We damaged the surface tissue of the ‘sphere’ with laser beams (using confocal microscopy), which released numerous bacteria from the ‘sphere’ (**Fig. 7**h-k), suggesting that these spherical formations encapsulate bacteria inside (**Fig. 7**g-g’’).

*Reversed nature of ‘spheres*. When we transferred 28 already separated alive ‘spheres’ (**S Fig. 5**) to new Petri dishes with algal mats, then within 3-5 days, we observed the restoration of classical placozoan bodyplan. During next few days, stable populations of placozoans could be established. In contrast, when we placed 28 control spheres in sterile Petri dishes without algal mats, all ‘spheres’ were degraded within 2-5 days.

*Fiber cells are capable of phagocytosis* (Theimann and Ruthman, 1990; Jacob et al., 2004; Smith and Reese, 2016; Moroz and Romanova, 2021) and contain bacterial cells (Guidi et al., 2011; Kamm et al., 2019; Gruber-Vodicka et al., 2019; Romanova et al., 2021). We also confirmed that in H1, H4, and H13 haplotypes, bacteria are localized only in fiber cells (**Fig. 8**b). Previously, it was hypothesized that bacteria could be endosymbionts. Moreover, the ultrastructural analysis of fiber cells suggested the engulfment of bacteria (**Fig. 7**b) by the endoplasmic reticulum, which can also be viewed as a stage of intracellular phagocytosis (**Fig. 8**b).

The treatment of *Trichoplax* with ampicillin eliminated bacteria from placozoans (Supplementary **Fig. 5S**) and their fiber cells (**Fig. 8f-i**). Furthermore, ampicillin prevented the formation of ‘spheres’ in bacteria-free culture: none were seen during 12 months of cultivating *Rickettsia*-free animals (over 10 000 animals, Supplementary **Fig. 5S**).

The fiber cell type contains the massive mitochondrial cluster (**Figs. 7**>b, **8**f-g; Grell, 1991; Smith et al., 2014; Mayorova et al., 2018; Romanova et al., 2021, Fig. 5) as a reporter of high energy production. Ampicillin-treated, *Rickettsia*-free, animals had additional morphologic features such as numerous small and clear inclusions inside fiber cells (**Fig. 8**f-g). The control group with bacteria has one or two large inclusions (**Fig. 8**h-i) with a less visible mitochondrial cluster. Functional significance of these ultrastructural changes is unclear.

## DISCUSSION

Grazing on algal and bacterial mats might be an ancestral feeding mode in early Precambrian animals (Rozhnov, 2009). And placozoans apparently preserved this evolutionarily conserved adaptation from Ediacaran animals (Sperling and Vinther, 2010). Under this scenario, we view the long-term culturing of placozoans as an essential paradigm to study interactions among relatively small numbers of cell types for integration of morphogenesis and reproduction, immunity and behaviors. Fig. 9 summarizes the complementary life-history strategies present in at least four haplotypes of Placozoa (H1, H2, H4, and H13).

**Figure 9.**
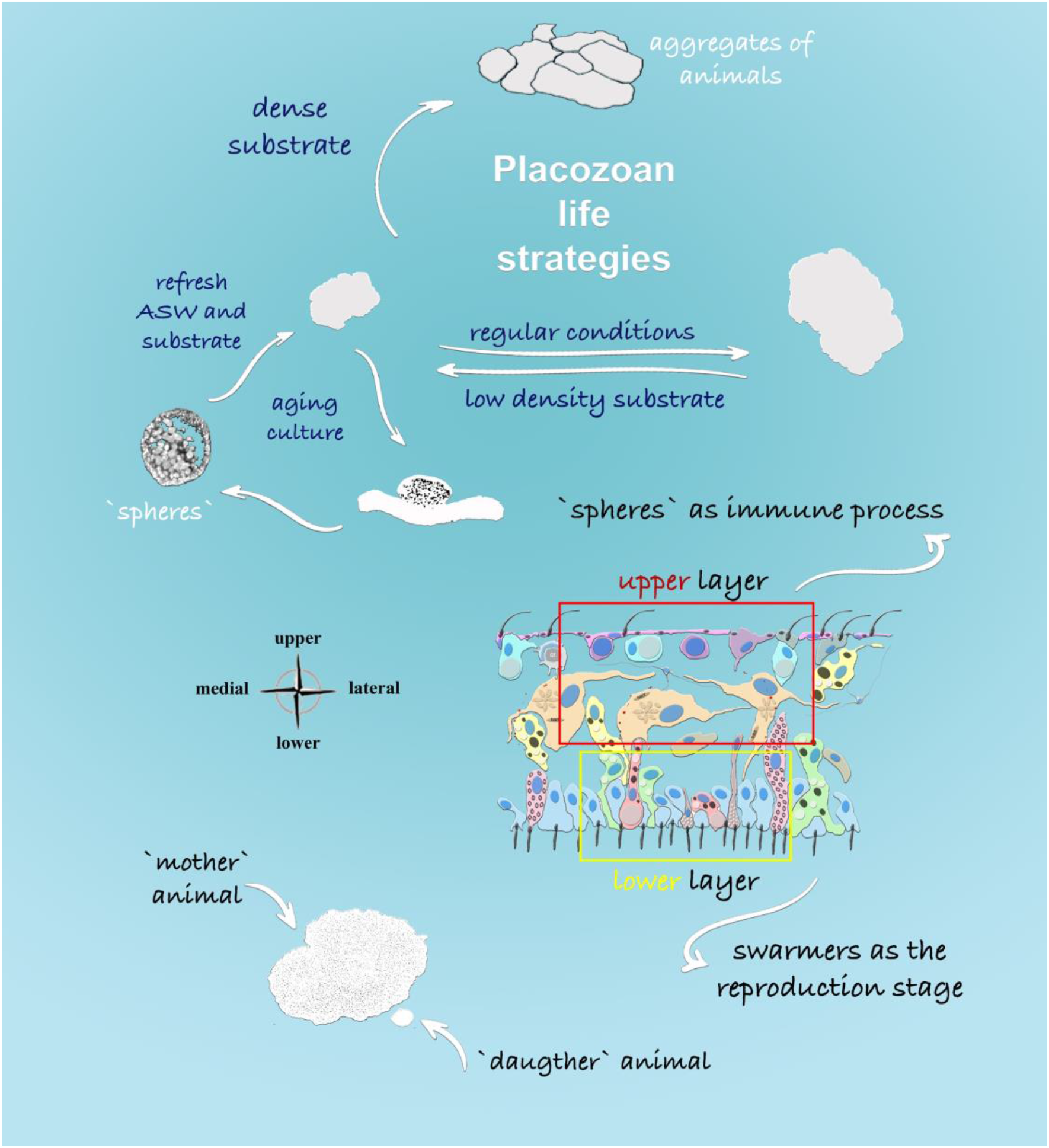
Life Strategies in Placozoa: Schematic representation of feeding and reproductive stages described in this study. The density of food substrate predominantly determines formations of different morphological stages. Dense algal substrate induced the formation of aggregates from multiple animals (‘social’ feeding behavior). In aging culture, the formation of specialized spherical structures was observed from the upper cell layer of placozoans, and it was shown that spheres could harbor multiple bacteria. In contrast, the formation of small juvenile animals or ‘swarmers’ might have different etiology and development from the lower cell layer, as shown in the cross-section of *Trichoplax*.

The directional (ciliated) locomotion in placozoans depends upon distributions of food sources, which differentially alternate exploratory and feeding patterns. Dense biofilms triggered social-type interactions and elementary cooperation, whereas limited food supplies, stress, and aging triggered systemic immune and morphogenic responses as well as alternative modes of reproduction.

Placozoans are virtually immortal with dominant clonal reproduction strategies (see Schierwater et al., 2021 and below). Furthermore, our data with small cell aggregates (section 2.2.) suggest that regenerative responses might also be part of adaptive reproduction mechanisms.

In addition, placozoans have two classical types of asexual reproduction: (i) fragmentation into two daughter animals or fission and (ii) formation of swarmers (Thiemann and Ruthmann, 1988, 1990, 1991; Seravin, 2004) – juvenile animals with small body size (20-30 µm). Thiemann and Ruthmann showed that the ‘budding’ of a daughter animal started from the dorsal side for 24 hours, and a released swarmer had all four morphologically defined cell types (Thiemann and Ruthmann, 1988, 1990, 1991; Seravin, 2004).

Here, we observed that the formation of juvenile animals could occur from the lower layer in all haplotypes of placozoans tested here. This type of arrangement might have some rationale: the lower epithelium consists of a greater diversity of cell types (compared to the upper layer) such as epithelial, lipophil, and gland cells with various subtypes (e.g., Smith et al., 2014; Mayorova et al, 2019; Romanova et al., 2021; Prakash et al., 2021).

The differences between the current and earlier observations on swarmers might be explained if we consider the formation of ‘spheres’ described here. But the spheres could be initially linked to the innate immune response, possibly induced by bacteria in aging populations or unhealthy culture conditions. Spheres, developed from the upper layer, could also be a path to asexual reproduction. If the microenvironment became more favorable for placozoans, ‘spheres’ can be similar (or transformed) to swarmers as juvenile animals. Thus, we think that in some earlier observations, physically separated spheres were described as swarmers.

One of the forms of nonspecific defense in invertebrates is the encapsulation of foreign objects. This process is similar to the systemic sphere formation in Placozoa. We see that opsonization has been observed inside the fiber cells (**Figs. 7-8**). Plus, the fiber cells can perform the functions of macrophages with a well-developed capacity for phagocytosis.

Cellular immunity by phagocytosis is the most ancient and widespread mechanism among basal Metazoa. For example, in sponges and cnidarians, the encapsulation is carried out by amoebocytes (=archeocytes) or collencytes (Musser et al., 2021). Phagocytosis in invertebrates, like in vertebrates, includes several stages: chemotaxis, recognition, attachment of a foreign agent to the phagocyte membrane, intracellular lysis, etc. (Bayne, 1990; Bang, 1975; Lackie, 1980). Due to the limited diversity of cell types, humoral and cellular immune responses could likely be relatively simple in Placozoa (Popgeorgiev et al., 2020).

Chemoattractant/repellents can be signaling molecules from microorganisms or other cell types. Symbiont-host signaling can include changes in nitric oxide gradients (Moroz et al., 2020b) or regional differences in amino acid composition, interconversion of D- and L-forms (Moroz et al., 2020a), the formation of oxygen radicals, which are toxic to bacteria, etc.

Fiber cells are perfectly suitable to be sensors, integrators, and effectors of the placozoan immune system. Fiber cells are located in the middle layer of placozoans with multiple elongated processes, spread around many other cell types, including the crystal cells. Fiber cells have specialized contacts among themselves (Grell and Ruthman, 1991). In the vicinity of fiber cells, a new class of neural-like/stellar-like cells are localized. Together they form a meshwork of cellular processes from the middle layer to the upper and lower layers (Moroz et al., 2021a; Romanova et al., 2021). This ‘network’ can be a functional integrative system sharing some immune and neural features and pools of signaling molecules in multiple microcavities for volume transmission (Moroz et al., 2021).

We propose that such placozoan-type integrative system is conceptually similar to the ancestral integration of immune and primordial neuroid-like systems, which controlled adaptive stress responses and behavior and regulated morphogenesis and regeneration.

Early (and present) animals strongly depended on the environmental and symbiotic bacteria, and fiber-type cells (or similar/homologous classes of amoebocytes as recently described in sponges - Musser et al, 2021) might be critical elements in the shared evolution of immune and neural systems to integrate both morphogenesis and behaviors (Fields et al., 2020). Here, the defense against bacterial infections can be an inherent part of such integrative ancestral adaptive responses.

## AUTHORS CONTRIBUTIONS

D.R. and L.L.M. designed the study; D.R. prepared illustrations; M.A.N. and D.R. cultured of all haplotypes; D.R. made microscopic and behavioral observations, and experiments of population growth rate; S.V.S. was involved in TE microscopy; M.A.N. designed and performed tests with antibiotics; D.R. and L.L.M. wrote the draft of the paper; and all authors reviewed, commented on, and edited the manuscript.

## ACKNOWLEDGMENTS

The authors thank Drs. M. Eitel and F. Varoqueaux for H2 and H13 haplotypes access, and A. Pronosin for his help in maintaining cultural dishes.

